# Cognition is associated with task-related brain network reconfiguration in late childhood

**DOI:** 10.1101/2025.03.26.645532

**Authors:** Mackenzie E Mitchell, Ashley J Jaimes, Tehila Nugiel

**Author notes:** Corresponding author: Tehila Nugiel, Department of Psychology, Florida State University, 1107 W Call St. Tallahassee, FL 32304.

## Abstract

In order to transition between a resting state and carrying out cognitively-demanding processes the brain makes a host of subtle changes to its network organization. In adults, less reconfiguration relates to better task performance, suggesting a preconfigured brain organization at rest is beneficial, such that only minute changes are required to execute task demands. Here, we take a developmental lens to this phenomenon, examining reconfiguration in late childhood by leveraging a large sample of 9-11 year olds from the Adolescent Brain and Cognitive Development Study. We find more reconfiguration between the resting state and two executive function tasks is related to better task performance, as well as better crystalized and fluid cognition in some cases. These relationships hold even when accounting for network segregation. These findings suggest a less-preconfigured, and thus more flexible, brain organization that enacts more reconfiguration to move from the resting state into a task state is beneficial in children. This aligns with theories positioning late childhood and the beginning of adolescence as a period of increased brain plasticity where functional brain networks are still undergoing refinement, and thus preconfiguration may be less beneficial and, instead, may place premature constraints on brain organization.

## 1. Introduction

One of the most striking properties of the brain is the ability to concurrently facilitate a variety of goals that change moment-to-moment and maintain a wealth of knowledge and skills that are stable throughout days, months, and years. In order to orchestrate this array of dynamic and stable functions, the brain simultaneously maintains a core functional organization and flexibly changes subtle aspects of its organization to facilitate ever changing task demands. These organizational properties can be attributed to large-scale functional brain networks (Bressler & Menon, 2010), characterized by distinct sets of brain regions that show stronger structural and functional connections to each other relative to other brain regions (Bassett & Sporns, 2017). This pattern of local coherence within networks paired with select connections between networks allows for both functional specialization (Buckner & Krienen, 2013) that supports local computations as well as cross-communication between distinct networks to give rise to multi-process complex behaviors (Dehaene et al., 1998; Sporns, 2013).

In adults, functional brain networks are highly stable both across people and within an individual (Gratton et al., 2018). However, there are small, but meaningful shifts in brain networks with changes in context, such as shifting from a resting state to a cognitive task, including tasks probing executive functions (Cohen & D’Esposito, 2016; Gratton et al., 2018; Krienen et al., 2014). This change in brain network organization from rest to task, which we will refer to as “reconfiguration,” has been shown to relate to state-based measures of cognition such as task performance (Cohen & D’Esposito, 2016; Schultz & Cole, 2016), as well as trait-based measures such as intelligence (Schultz & Cole, 2016). Specifically, less reconfiguration, or more similarity between rest and task states, is related to higher intelligence in adults (Schultz & Cole, 2016). This work suggests that optimal cognition is promoted by a “preconfigured” intrinsic brain network organization that facilitates an efficient transition into a task state.

Functional brain networks show rapid organization in early life and then slowly refine into adulthood (Fransson et al., 2007, 2010; Gilmore et al., 2018; Grayson & Fair, 2017). Sensorimotor networks exhibit rapid development with significant maturation across the first two years of life (Cao et al., 2017; Gilmore et al., 2018). Brain networks that support higher-order processes and facilitate communication and information integration across networks are slower to mature (Grayson & Fair, 2017; Larsen et al., 2023) and show significant change and refinement throughout adolescence (Cui et al., 2020). During this period of refinement, functional networks and their boundaries are not yet clearly defined (Tooley et al., 2022), which suggests a potential lack of specialization. As networks are poorly defined in youth, it is unclear whether preconfiguration of brain network organization would be efficient, or is even established, in the developing brain. Previous work in late childhood suggests that the similarity between brain network organization at rest and during tasks is also linked to task performance (Le et al., 2020), although findings vary depending on the task type (e.g. working memory vs. perceptual processing). This distinction in task type could speak to an important developmental nuance where preconfiguration is beneficial for some domains of cognition but not others. An understanding of how preconfiguration changes with age and in what contexts it is beneficial can be useful for understanding how brain network maturation and refinement supports the cognitive maturation seen across adolescence (Tervo-Clemmens et al., 2022).

The current study uses a sample of thousands of youth in late childhood from the Adolescent Brain and Cognitive Development (ABCD) Study (Casey et al., 2018) to examine how the brain reconfigures between rest and task states. Using two cognitively-demanding but distinct tasks tapping working memory and response inhibition, we quantified the degree of brain network reconfiguration between the resting and each task state. We tested whether reconfiguration predicts task performance and broad measures of general cognition. It is possible that, similar to adults, preconfigured network organization in youth is related to better cognition. It is alternatively possible that in youth a preconfigured network organization could be counterproductive, restricting plasticity and the opportunity for experience to continue to refine brain network organization late into adolescence. This study will clarify between these options and allow us to better understand where youth between the ages of 9 and 11 years lie in the spectrum between flexibility and preconfiguration and how individual differences in this maturational trajectory predict cognition. We will further explore how the degree of reconfiguration relates to intrinsic brain network organization and differences across age.

## 2. Methods

### 2.1. Sample and demographics

This project utilized data from 11,875 youth aged 9-10 years in the baseline collection of the ABCD Study (Release 3.0; doi: 10.15154/1520591) (Casey et al., 2018). Functional MRI data was acquired from the ABCD-BIDS Collection 3165 (Feczko et al., 2021). Parents provided written informed consent and participants provided assent prior to participation in the study. For inclusion in this project, participants were required to have high-quality fMRI data in the resting state and in at least one of the executive function tasks (SST, EN-back), as well as behavioral performance meeting ABCD quality control criteria in the corresponding tasks. After excluding participants who did not meet these criteria, the final sample in this project consisted of *N* = 4,941 participants.

Parents of participants reported the following demographics on each participant: age, sex, race/ethnicity, parental education, family income, and parental marital status (see **Table 1**). An index of neighborhood disadvantage was calculated based on each participant’s home address (Fan et al., 2021; Kind et al., 2014) and provided by the ABCD Study. We constructed a composite socioeconomic status score from a confirmatory factor analysis of family income, parental education in years, and neighborhood disadvantage index, as in (Rakesh et al., 2021; Sripada et al., 2021). We converted family income from categorical bins (e.g., “$5,000 through $11,999”, “$75,000 through $99,999”) to numeric quantities by taking the natural log of the highest value of the lowest bin (i.e., “Less than $5,000”: $5,000), the lowest value from the highest bin (i.e., “$200,000 and greater”: $200,000), and the midpoint from every other bin (e.g., “$75,000 through $99,999”: $87,499.50). We also converted the highest level of parental education from categories(e.g., “GED or equivalent”) into a numeric value representing years of education (e.g., “GED or equivalent”: 12 years).

**Table 1.**
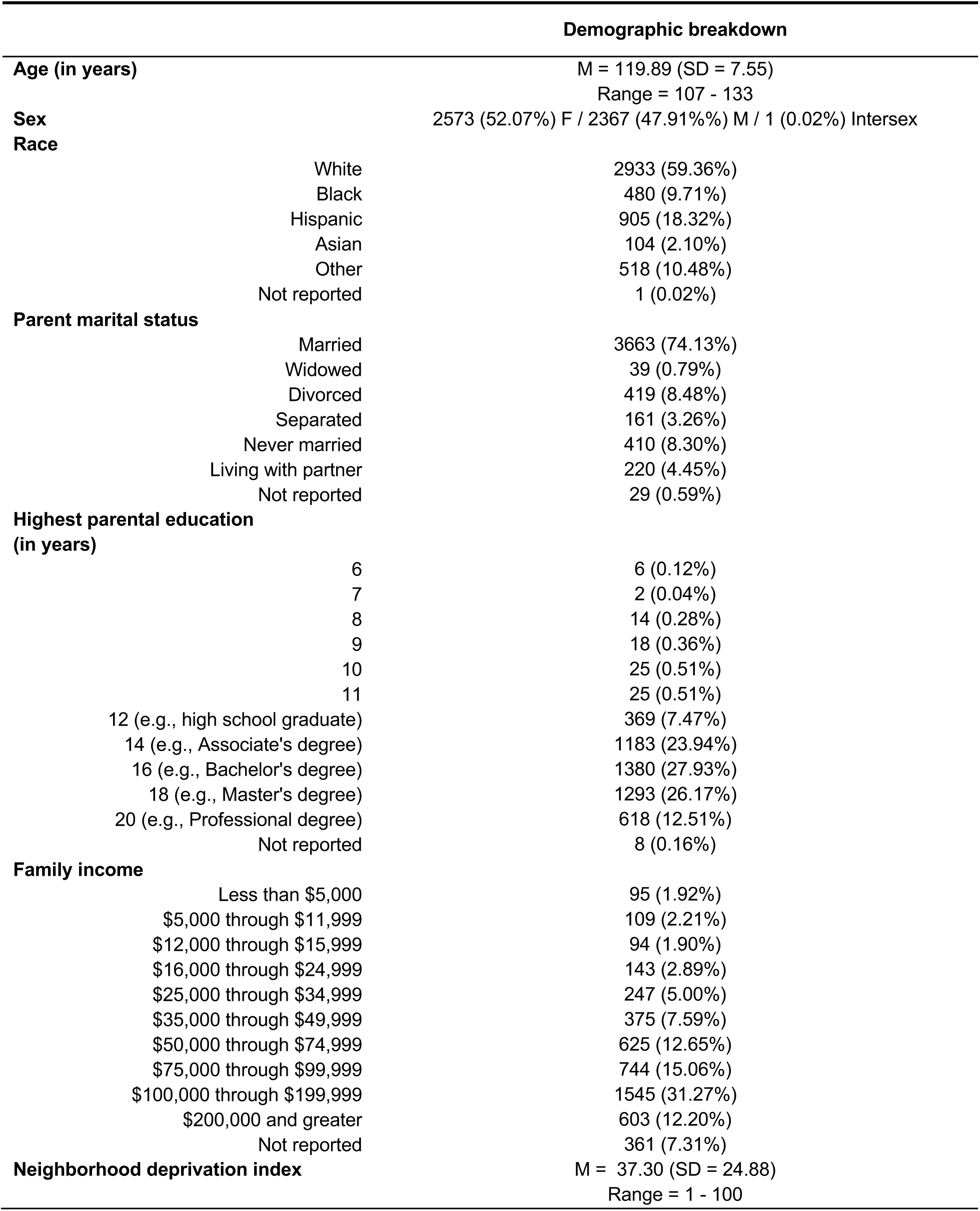
Demographic characteristics.

|  |  | Demographic breakdown |
| --- | --- | --- |
| **Age (in years)** |  | M = 119.89 (SD = 7.55)<br>Range = 107 - 133 |
| **Sex** |  | 2573 (52.07%) F / 2367 (47.91%) M / 1 (0.02%) Intersex |
| **Race** |  |  |
|  | White | 2933 (59.36%) |
|  | Black | 480 (9.71%) |
|  | Hispanic | 905 (18.32%) |
|  | Asian | 104 (2.10%) |
|  | Other | 518 (10.48%) |
|  | Not reported | 1 (0.02%) |
| **Parent marital status** |  |  |
|  | Married | 3663 (74.13%) |
|  | Widowed | 39 (0.79%) |
|  | Divorced | 419 (8.48%) |
|  | Separated | 161 (3.26%) |
|  | Never married | 410 (8.30%) |
|  | Living with partner | 220 (4.45%) |
|  | Not reported | 29 (0.59%) |
| **Highest parental education (in years)** |  |  |
|  | 6 | 6 (0.12%) |
|  | 7 | 2 (0.04%) |
|  | 8 | 14 (0.28%) |
|  | 9 | 18 (0.36%) |
|  | 10 | 25 (0.51%) |
|  | 11 | 25 (0.51%) |
|  | 12 (e.g., high school graduate) | 369 (7.47%) |
|  | 14 (e.g., Associate's degree) | 1183 (23.94%) |
|  | 16 (e.g., Bachelor's degree) | 1380 (27.93%) |
|  | 18 (e.g., Master's degree) | 1293 (26.17%) |
|  | 20 (e.g., Professional degree) | 618 (12.51%) |
|  | Not reported | 8 (0.16%) |
| **Family income** |  |  |
| | Less than \$5,000 | 95 (1.92%) |
| | \$5,000 through \$11,999 | 109 (2.21%) |
| | \$12,000 through \$15,999 | 94 (1.90%) |
| | \$16,000 through \$24,999 | 143 (2.89%) |
| | \$25,000 through \$34,999 | 247 (5.00%) |
| | \$35,000 through \$49,999 | 375 (7.59%) |
| | \$50,000 through \$74,999 | 625 (12.65%) |
| | \$75,000 through \$99,999 | 744 (15.06%) |
| | \$100,000 through \$199,999 | 1545 (31.27%) |
| | \$200,000 and greater | 603 (12.20%) |
|  | Not reported | 361 (7.31%) |
| **Neighborhood deprivation index** |  | M = 37.30 (SD = 24.88)<br>Range = 1 - 100 |

### 2.2. Brain data descriptions

Participants underwent a functional MRI session, including runs of resting state, an emotional n-back (EN-back) task, and a stop signal task (SST). See (Casey et al., 2018) for a full description of the ABCD Study neuroimaging protocol and fMRI tasks; we have included relevant information for the current analyses here.

During the resting state, participants were instructed to stay awake and passively view a cross hair. Participants completed up to four runs of resting state, each five minutes in duration.

During the EN-back task probing working memory (Barch et al., 2013; Casey et al., 2018), participants viewed a series of images arranged in blocks by type: places, happy faces, neutral faces, fearful faces. On some blocks, participants were tasked to report if each image matched or did not match the target image shown at the beginning of the block, termed a “0-back block.” On other blocks, participants were tasked to report if each image matched or did not match the image shown two prior, termed a “2-back block.” Participants completed up to two runs of the EN-back task, each 3 minutes and 44 seconds in duration. Each run contained four 2-back blocks (one for each image type; 10 trials each) and four 0-back blocks (one for each stimulus type 10 trials each). For this project, only performance and brain function on the 2-back blocks, which probe a higher-load of working memory than the 0-back blocks, were used. See Casey et al. (2018) for details on task timing.

During the SST probing response inhibition (Casey et al., 2018; Logan & Cowan, 1984), participants pressed buttons to report the direction of horizontal arrows (i.e., left or right) presented on the screen (“go” trials). Occasionally, the horizontal arrow would be interrupted by a vertical arrow, termed the “stop signal” (“stop” trials). Participants were instructed to suppress their button press on these stop trials. A staircase procedure was used to adjust the presentation time of subsequent stop signals. Specifically, when a participant failed to stop the button press at the presentation of a stop signal, the next stop signal would be presented 50ms earlier. When a participant succeeded in stopping the button press, the next stop signal would be presented 50ms later. By the end of each run, there should be approximately 50% successful and 50% unsuccessful stop trials. Participants completed up to two runs of the SST, each 5 minutes and 49 seconds in duration. Each run contained 180 trials (150 go trials and 30 stop trials). See Casey et al. (2018) for details on task timing.

### 2.3. Brain data collection and processing

Neuroimaging data were downloaded from the ABCD-BIDS Collection 3165 v1.1.1 from the NIMH Data Archive (NDA) (https://collection3165.readthedocs.io/en/stable/) (Feczko et al., 2021). Briefly, neuroimaging data were collected on three 3T scanner platforms (Siemens Prisma, General Electric 750, and Philips) with multi-channel head coils at 21 ABCD Study sites in the United States. Scanner acquisition parameters were harmonized across the three scanner platforms; see (Casey et al., 2018) for full details. High-resolution anatomical images were collected using 3D T1-weighted and T2-weighted images. Functional data were collected with a T2-weighted multiband EPI sequence with a slice acceleration factor of 6 (60 slices, 90x90 matrix, TR = 800 ms, TE = 30 ms, field of view 216 x 216mm, voxel dimensions: 2.4 mm x 2.4 mm x 2.4 mm). Each MRI session included a series of functional resting state and task runs, counterbalanced across participants.

Neuroimaging data were processed using the ABCD-BIDS pipeline (https://github.com/ABCD-STUDY/abcd-hcp-pipeline) (Fair et al., 2020; Feczko et al., 2021; Miranda-Domínguez et al., 2020). Initial processing of functional MRI data included distortion correction with spin echo field maps, intensity normalization, and registration of functional images to the native T1 and the MNI template. The functional data were projected onto individualized cortical surfaces created from anatomical scans using Freesurfer (Reuter et al., 2010) and then smoothed with a 2mm full-width-half-maximum kernel. Additional steps were taken to reduce the impact of head motion and respiration on the functional data. First, the time series were detrended and normalized. Then, the data underwent nuisance regression with six motion parameters, mean time series for white matter, cerebrospinal fluid, and global signal, as well as derivatives and squares for each. Next, a band-pass filter between 0.008 and 0.1 Hz using a 2nd order Butterworth filter (Hallquist et al., 2013) was applied along with spectral interpolation of frames with a filtered FD > 0.3mm. Then, a temporal notch filter between 0.31-0.43Hz was implemented to attenuate respiration artifacts in the timeseries motion estimates (Fair et al., 2020). Lastly, all functional runs in each cognitive state were concatenated, high motion timepoints with a filtered FD > 0.2mm were removed (“scrubbed”) from the time series (Power et al., 2012), and each concatenated timeseries was truncated to be the same length across all subjects (i.e., 5 minutes (240 timepoints) for the resting state and the SST; 2 minutes (96 timepoints) for the EN-back task 2-back blocks only). The concatenated data in each cognitive state was parcellated into regions using the Gordon cortical parcellation (Gordon et al., 2016), comprised of 333 cortical surface regions across 12 functional networks and an unassigned network.

### 2.4. Functional connectivity calculation and matrix formation

Pearson’s correlations were computed between each pair of regions to estimate functional connectivity in the resting state, EN-back task, and SST separately. This resulted in a 333 x 333 correlation matrix per cognitive state per participant. Each functional connectivity matrix was Fisher z-transformed prior to analyses. Each region was assigned to a brain network based on the Gordon parcellation (Gordon et al., 2016).

### 2.5. Reconfiguration measured with functional connectivity similarity between cognitive states

Reconfiguration between the resting state and each executive function task (EN-back, SST) was operationalized by measuring the similarity of functional connectivity matrices for each participant across pairs of cognitive states. For each participant and each cognitive state comparison, whole-brain functional connectivity similarity was calculated by vectorizing the lower triangle of the unthresholded, Fisher z-transformed correlation matrices for each cognitive state and correlating the pair of vectors (i.e., resting state and EN-back, resting state and SST), as in (Schultz & Cole, 2016). Within-participant functional connectivity similarity was also calculated for each large-scale brain network (except the unassigned network) by creating a vector of the connections within each network and between that network with every other network. This resulted in 13 measures of within-participant functional connectivity similarity correlations for each comparison (one whole brain and 12 networks), which were Fisher z-transformed to allow for further statistical analyses, as in (Schultz & Cole, 2016). Importantly, greater functional connectivity similarity values denote less reconfiguration between the resting state and each task.

See the *Supplementary Materials* for differences in functional connectivity similarity between the resting state and the EN-back task and the resting state and the SST.

### 2.6. Brain network organization

To probe how reconfiguration between the resting state and each executive function task related to resting-state brain network organization, we calculated global system segregation (Chan et al., 2014) during the resting-state as the difference between average within-network connectivity and average between-network connectivity divided by the average within-network connectivity. Global system segregation has been found to increase across childhood and adolescence (Keller et al., 2023; Sun et al., 2024; Tooley, Park, et al., 2022).

### 2.7. Cognitive measures of interest

We were first interested in how the degree of reconfiguration between the resting state and tasks related to cognitive abilities related probed in the executive function tasks of interest. To index working memory during the EN-back task, we used average percent accuracy on the 2-back blocks across both runs (M = 0.81, SD = 0.08, range = 0.61 - 0.99). To index response inhibition during the SST, we used mean stop signal reaction time (SSRT), or the mean reaction time on correct go trials minus the mean SSD, averaged across both runs (M = 294.09, SD = 61.40, range = 32.20 - 556.91).

We were also interested in how the degree of reconfiguration between the resting state and tasks related to broad measures of cognition. We used the uncorrected fluid and crystallized cognition composite scores from the NIH Toolbox battery (Akshoomoff et al., 2013). The fluid cognition composite is an average of standard scores on the pattern comparison processing speed, list sorting working memory, picture sequence memory, flanker, and dimensional change card sort tests (M = 93.87, SD = 9.74, range = 52 - 122). The crystallized cognition composite is an average of standard scores on the picture vocabulary and oral reading recognition tests (M = 87.68, SD = 6.56, range = 51 - 115).

### 2.8. Analyses

First, we tested for relationships between functional connectivity similarity and the measures of task performance on the fMRI executive function tasks (i.e., 2-back accuracy on the EN-back task, SSRT on the SST). To do this we estimated linear regressions using the *lme4* and *lmerTest* R packages (Kuznetsova et al., 2017) separately testing for relationships between 2-back accuracy and each measure of functional connectivity between the resting state and the EN-back task, as well as between the SSRT and each measure of functional connectivity between the resting state and the SST. We corrected for multiple comparisons among the p-values for the 12 network models for each functional connectivity similarity comparison (2 corrections with 12 p-values each) using a Benjamini-Hochberg false discovery rate (FDR) correction at q < .05 (Benjamini & Hochberg, 1995).

Then, we tested for relationships between functional connectivity similarity and broad measures of cognitive ability. To do this we estimated linear regressions separately testing for relationships between fluid cognition composite and each measure of functional connectivity similarity, as well as crystallized cognition composite and each measure of functional connectivity similarity. We corrected for multiple comparisons among the p-values for the 12 network models for each task and broad cognitive measure (4 corrections with 12 p-values each) using a FDR correction at q < .05.

Next, we tested how functional connectivity similarity relates to age. To do this we estimated a series of linear regressions separately with each functional connectivity similarity measure as the outcome and age as the predictor. We corrected for multiple comparisons among the p-values for the 12 network models for each functional connectivity similarity comparison (2 corrections with 12 p-values each) using a FDR correction at q < .05.

Finally, we tested how functional connectivity similarity relates to resting state brain network organization (global system segregation). Using linear regressions, we investigated relationships between global system segregation and each functional connectivity similarity measure. We corrected for multiple comparisons among the p-values for the 12 network models for each functional connectivity similarity comparison (2 corrections with 12 p-values each) using a FDR correction at q < .05 . Additionally, to understand if the relationships between functional connectivity and cognition could be driven by global system segregation, we probed how the global system segregation related to age, SSRT and 2-back accuracy, and fluid and crystallized cognition composites. Further, we ran additional linear regressions to test if functional connectivity similarity remained a significant predictor of cognitive measures when covarying for global system segregation. We corrected for multiple comparisons across all p-values using a FDR correction at q < .05.

All models included covariates of age, sex, socioeconomic status composite score, race/ethnicity, and parent marital status.

## 3. Results

### 3.1. Relationships between functional connectivity similarity and executive function task performance

First, we tested the similarity of functional connectivity between the resting state and each executive function task related to performance on the respective executive function task (**Figure 1**). We found that less whole brain similarity between the resting state and the EN-back task related to greater 2-back accuracy, or better performance on the EN-back task (*β* = -0.07, *SE* = 0.03, *p* < .006). Further, less similarity of the cingulo-opercular (*β* = -0.14, *SE* = 0.02, adjusted-*p* < .001), salience (*β* = -0.08, *SE* = 0.02, adjusted-*p* = .001), and dorsal somatomotor (*β* = -0.10, *SE* = 0.02, adjusted-*p* < .001) networks between the resting state and the EN-back task related to better 2-back accuracy. All other *p* values > .057 (**Table 2**).

**Table 2.**
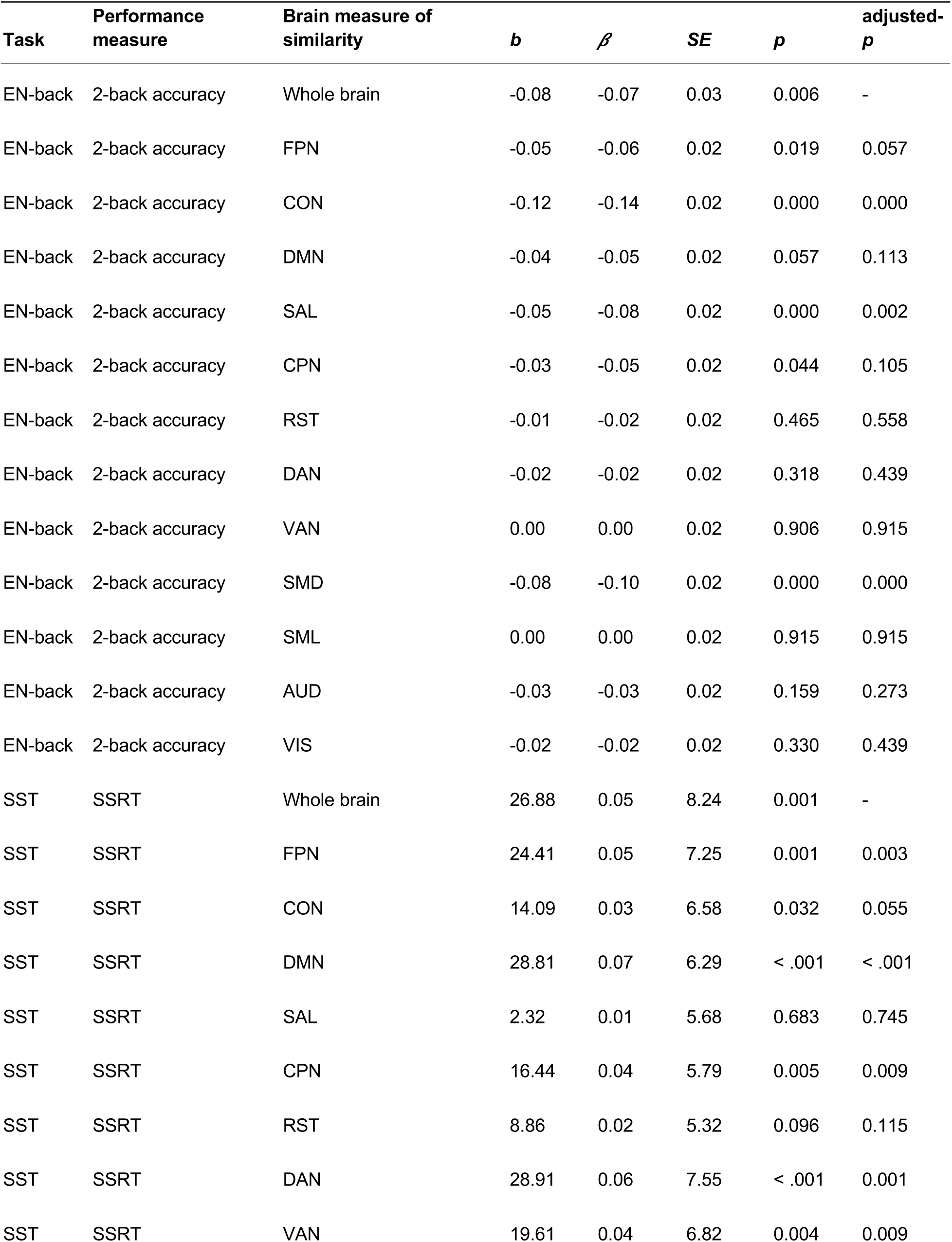
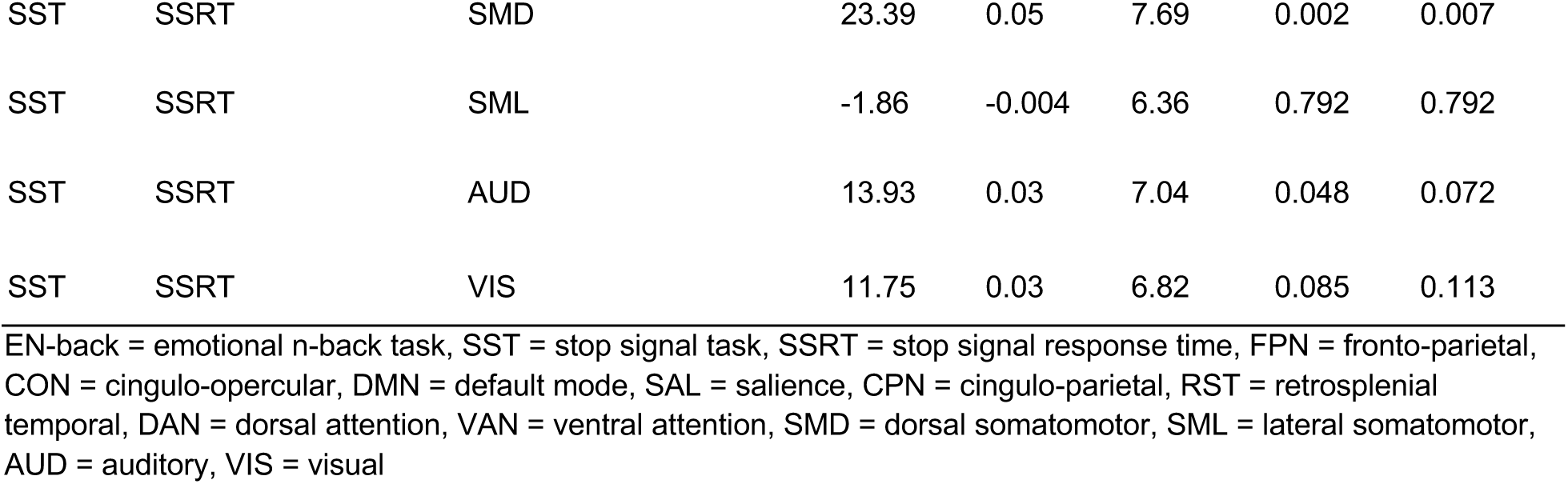
Functional connectivity similarity related to executive function task performance.

| Task | Performance measure | Brain measure of similarity | *b* | $\beta$ | *SE* | *p* | adjusted-*p* |
| --- | --- | --- | --- | --- | --- | --- | --- |
| EN-back | 2-back accuracy | Whole brain | -0.08 | -0.07 | 0.03 | 0.006 | - |
| EN-back | 2-back accuracy | FPN | -0.05 | -0.06 | 0.02 | 0.019 | 0.057 |
| EN-back | 2-back accuracy | CON | -0.12 | -0.14 | 0.02 | 0.000 | 0.000 |
| EN-back | 2-back accuracy | DMN | -0.04 | -0.05 | 0.02 | 0.057 | 0.113 |
| EN-back | 2-back accuracy | SAL | -0.05 | -0.08 | 0.02 | 0.000 | 0.002 |
| EN-back | 2-back accuracy | CPN | -0.03 | -0.05 | 0.02 | 0.044 | 0.105 |
| EN-back | 2-back accuracy | RST | -0.01 | -0.02 | 0.02 | 0.465 | 0.558 |
| EN-back | 2-back accuracy | DAN | -0.02 | -0.02 | 0.02 | 0.318 | 0.439 |
| EN-back | 2-back accuracy | VAN | 0.00 | 0.00 | 0.02 | 0.906 | 0.915 |
| EN-back | 2-back accuracy | SMD | -0.08 | -0.10 | 0.02 | 0.000 | 0.000 |
| EN-back | 2-back accuracy | SML | 0.00 | 0.00 | 0.02 | 0.915 | 0.915 |
| EN-back | 2-back accuracy | AUD | -0.03 | -0.03 | 0.02 | 0.159 | 0.273 |
| EN-back | 2-back accuracy | VIS | -0.02 | -0.02 | 0.02 | 0.330 | 0.439 |
| SST | SSRT | Whole brain | 26.88 | 0.05 | 8.24 | 0.001 | - |
| SST | SSRT | FPN | 24.41 | 0.05 | 7.25 | 0.001 | 0.003 |
| SST | SSRT | CON | 14.09 | 0.03 | 6.58 | 0.032 | 0.055 |
| SST | SSRT | DMN | 28.81 | 0.07 | 6.29 | < .001 | < .001 |
| SST | SSRT | SAL | 2.32 | 0.01 | 5.68 | 0.683 | 0.745 |
| SST | SSRT | CPN | 16.44 | 0.04 | 5.79 | 0.005 | 0.009 |
| SST | SSRT | RST | 8.86 | 0.02 | 5.32 | 0.096 | 0.115 |
| SST | SSRT | DAN | 28.91 | 0.06 | 7.55 | < .001 | 0.001 |
| SST | SSRT | VAN | 19.61 | 0.04 | 6.82 | 0.004 | 0.009 |
| SST | SSRT | SMD | 23.39 | 0.05 | 7.69 | 0.002 | 0.007 |
| SST | SSRT | SML | -1.86 | -0.004 | 6.36 | 0.792 | 0.792 |
| SST | SSRT | AUD | 13.93 | 0.03 | 7.04 | 0.048 | 0.072 |
| SST | SSRT | VIS | 11.75 | 0.03 | 6.82 | 0.085 | 0.113 |

1 EN-back = emotional n-back task, SST = stop signal task, SSRT = stop signal response time, FPN = fronto-parietal, CON = cingulo-opercular, DMN = default mode, SAL = salience, CPN = cingulo-parietal, RST = retrosplenial temporal, DAN = dorsal attention, VAN = ventral attention, SMD = dorsal somatomotor, SML = lateral somatomotor, AUD = auditory, VIS = visual

We also found that less whole brain similarity between the resting state and the SST related to faster SSRT, or better performance on the SST (*β* = 0.05, *SE* = 8.24, adjusted-*p* = .001). Further, less similarity of the fronto-parietal (*β* = 0.05, *SE* = 7.25, adjusted-*p* = .003), dorsal attention (*β* = 0.06, *SE* = 7.55, adjusted-*p* < .001), ventral attention (*β* = 0.04, *SE* = 6.82, adjusted-*p* = .009), default mode (*β* = 0.07, *SE* = 6.29, adjusted-*p* < .001), cingulo-parietal (*β* = 0.04, *SE* = 5.79, adjusted-*p* = .009), and dorsal somatomotor (*β* = 0.05, *SE* = 7.69, adjusted-*p* = .007) networks between the resting state and the SST related to lower SSRT. Together, across both executive function tasks, less functional connectivity similarity to, or greater reconfiguration from, the resting state is related to better task performance, indexed by higher 2-back accuracy and lower SSRT. All other *p* values > .055 (**Table 2**).

**Figure 1.**
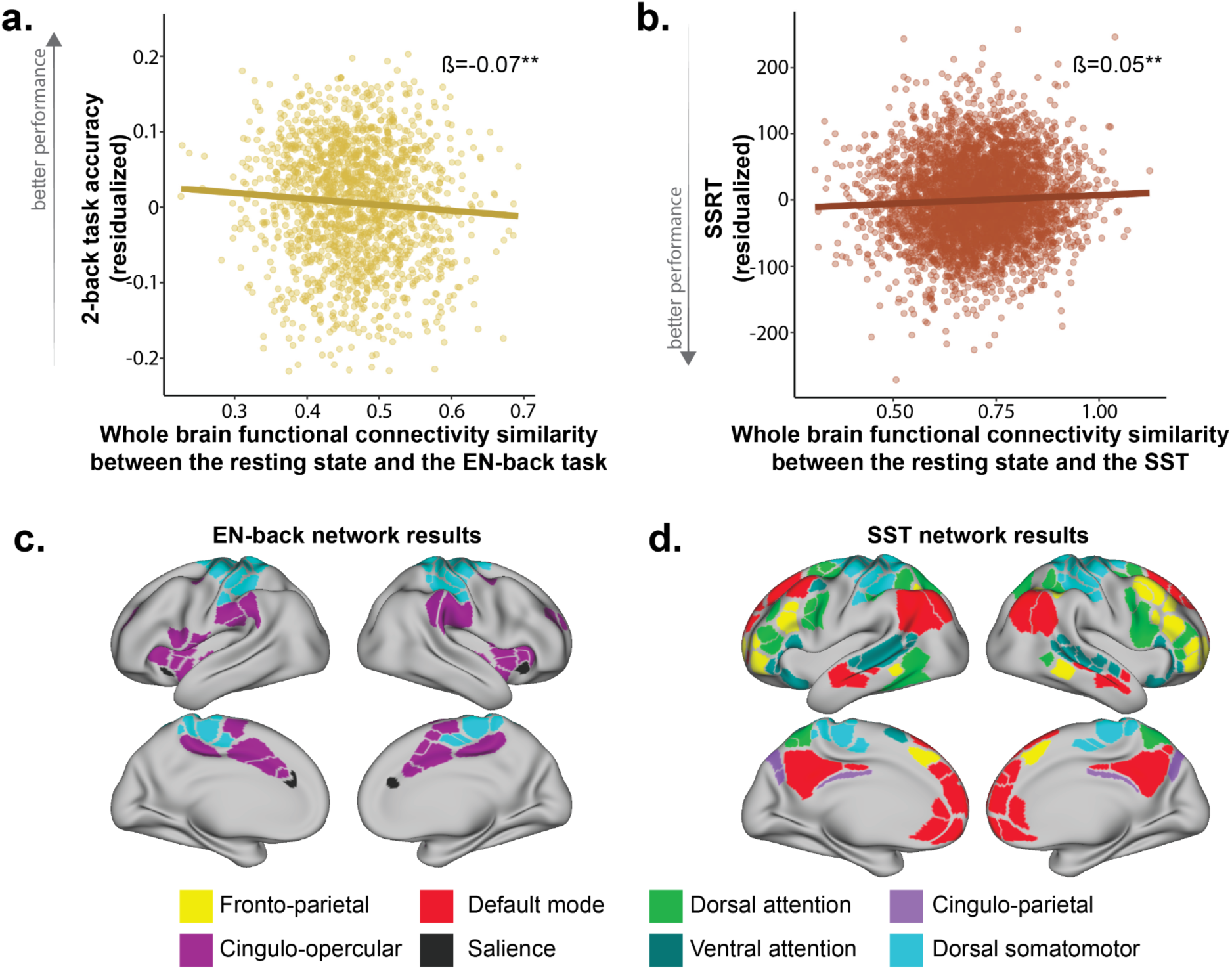
Functional connectivity similarity to the resting state is related to executive function task performance. **a.** Lower whole brain functional connectivity similarity between the resting state and the EN-back task (yellow) is related to greater 2-back task accuracy. **b.** Lower whole brain functional connectivity similarity between the resting state and the SST (red) is related to faster SSRT, or better performance. **c.** Functional connectivity similarity of the cingulo-opercular (dark purple), salience (black), and dorsal somatomotor (light blue) networks between the resting state and the EN-back tasks are related to 2-back task accuracy. **d.** Functional connectivity similarity of the fronto-parietal (yellow), cingulo-opercular (dark purple), default mode (red), cingulo-opercular (light purple), dorsal attention (green), ventral attention (teal), and dorsal somatomotor (light blue) networks between the resting state and the SST are related to SSRT. EN-back = Emotional n-back task; SST = Stop signal task; SSRT = Stop signal response time; \**p*<.05; \*\**p*<.01; \*\*\**p*<.001

### 3.2. Relationships between functional connectivity similarity and broad cognitive performance

To understand if the relationship between functional connectivity similarity and cognition extends beyond performance on the examined fMRI executive function tasks, we probed relationships with measures of broad cognitive performance from the NIH Toolbox (**Figure 2**). We found that less whole brain functional connectivity similarity between the resting state and the EN-back task related to greater fluid cognition (*β* = -0.05, *SE* = 3.11, *p* = .042) and greater crystallized cognition (*β* = -0.06, *SE* = 1.96, *p* = .011) on the NIH Toolbox battery. Additionally, less cingulo-opercular functional connectivity similarity between the resting state and the EN-back task related to greater crystallized cognition (*β* = -0.07, *SE* = 1.41, adjusted-*p* = .021). All other adjusted-*p* values > .107 (**Table 3**).

**Figure 2.**
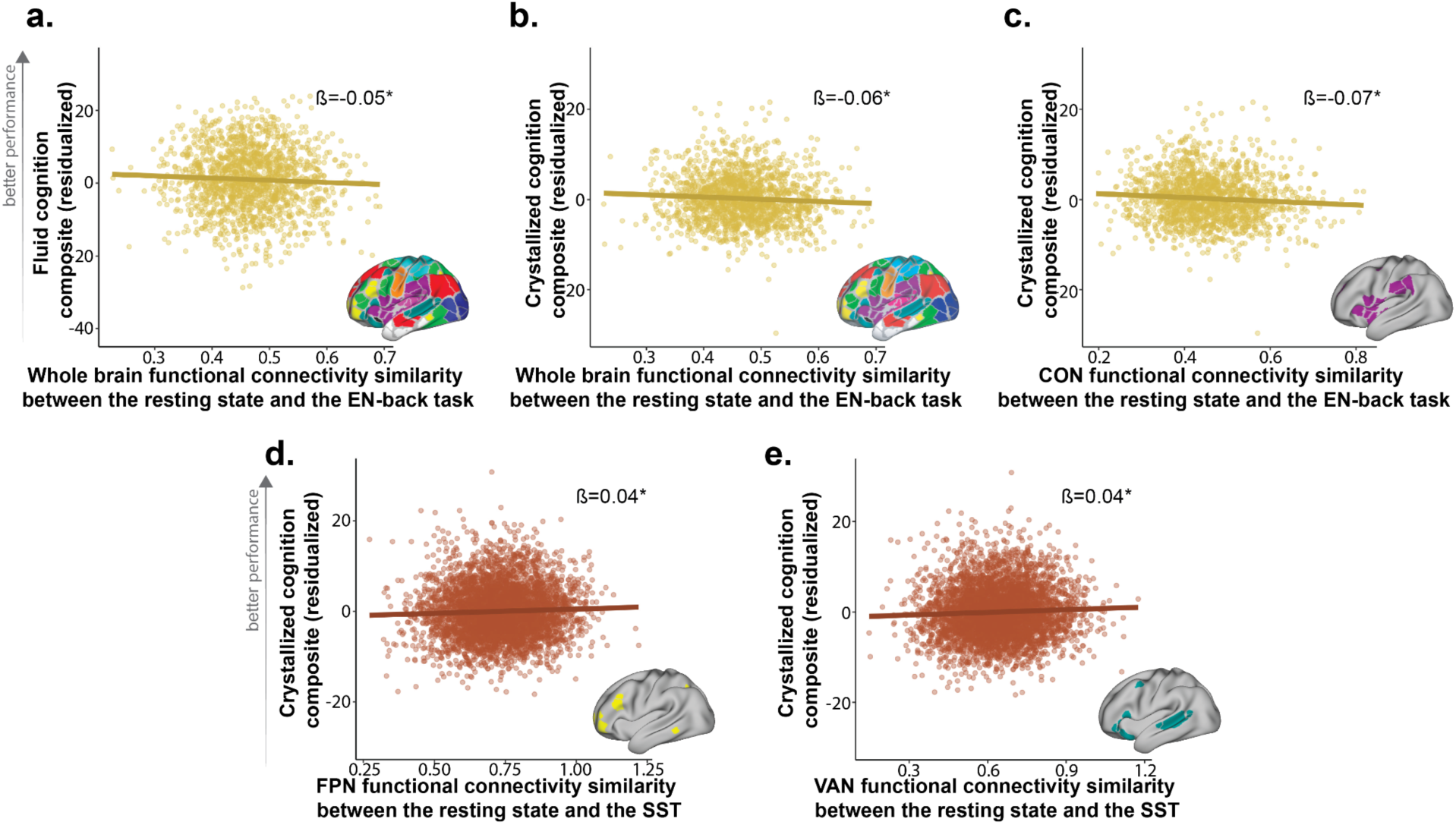
Functional connectivity similarity to the resting state is related to broad measures of cognition. Less whole brain functional connectivity similarity between the resting state and the EN-back task (yellow) is related to better **(a)** fluid cognition and **(b)** crystallized cognition. **(c)** Less functional connectivity similarity of the cingulo-opercular network between the resting state and the EN-back task (yellow) is related to better crystallized cognition. Greater functional connectivity similarity of the **(d)** fronto-parietal and **(e)** ventral attention networks between the resting state and the SST (red) are related to better crystallized cognition. EN-back = Emotional n-back task; SST = Stop signal task; *(adjusted-)*p*<.05; **(adjusted-)*p*<.01; ***(adjusted-)*p*<.001

**Table 3.**
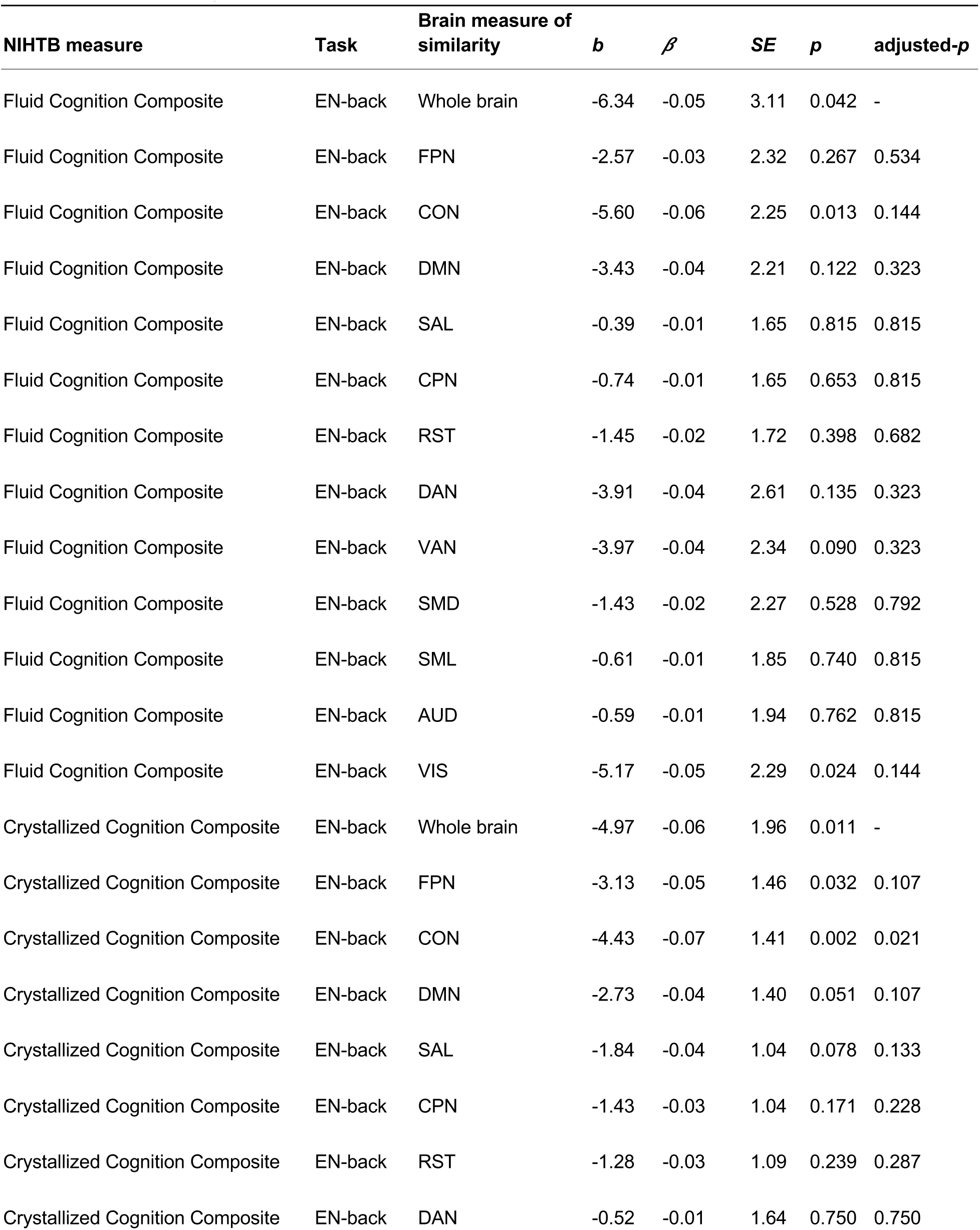
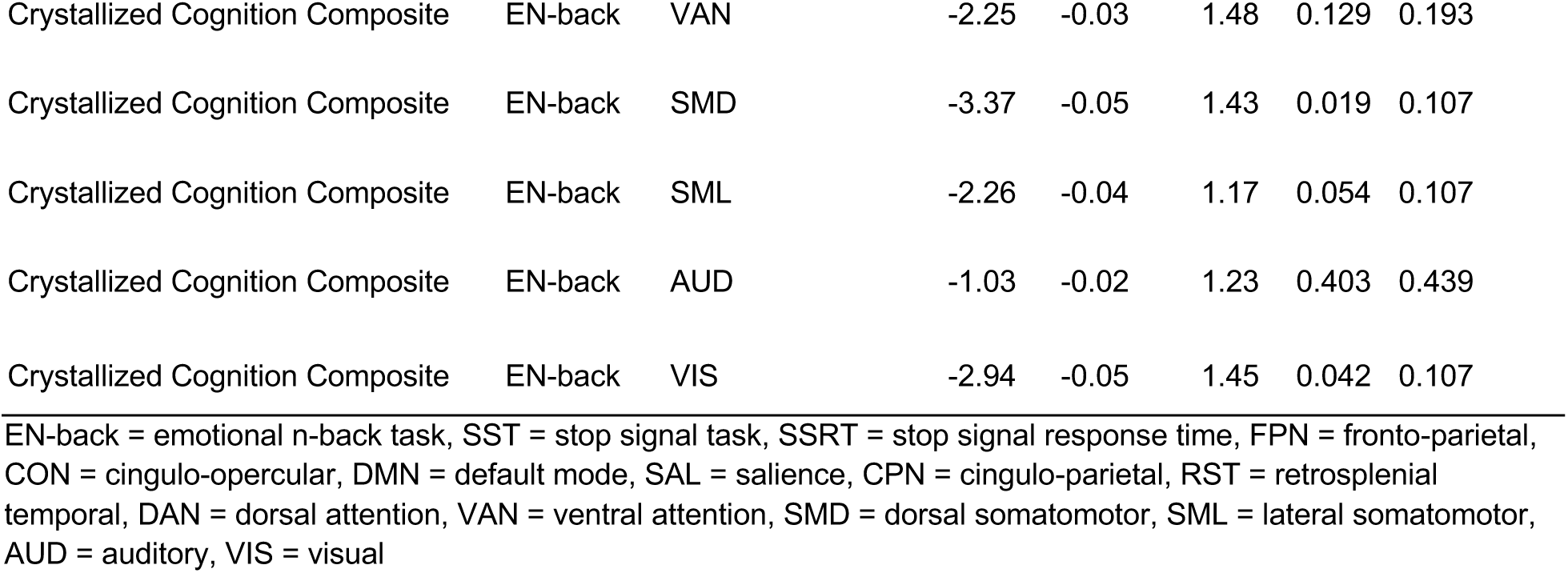
Functional connectivity similarity between the resting state and EN-back task related to broad cognitive measures.

| NIHTB measure | Task | Brain measure of similarity | *b* | $\beta$ | *SE* | *p* | adjusted-*p* |
| --- | --- | --- | --- | --- | --- | --- | --- |
| Fluid Cognition Composite | EN-back | Whole brain | -6.34 | -0.05 | 3.11 | 0.042 | - |
| Fluid Cognition Composite | EN-back | FPN | -2.57 | -0.03 | 2.32 | 0.267 | 0.534 |
| Fluid Cognition Composite | EN-back | CON | -5.60 | -0.06 | 2.25 | 0.013 | 0.144 |
| Fluid Cognition Composite | EN-back | DMN | -3.43 | -0.04 | 2.21 | 0.122 | 0.323 |
| Fluid Cognition Composite | EN-back | SAL | -0.39 | -0.01 | 1.65 | 0.815 | 0.815 |
| Fluid Cognition Composite | EN-back | CPN | -0.74 | -0.01 | 1.65 | 0.653 | 0.815 |
| Fluid Cognition Composite | EN-back | RST | -1.45 | -0.02 | 1.72 | 0.398 | 0.682 |
| Fluid Cognition Composite | EN-back | DAN | -3.91 | -0.04 | 2.61 | 0.135 | 0.323 |
| Fluid Cognition Composite | EN-back | VAN | -3.97 | -0.04 | 2.34 | 0.090 | 0.323 |
| Fluid Cognition Composite | EN-back | SMD | -1.43 | -0.02 | 2.27 | 0.528 | 0.792 |
| Fluid Cognition Composite | EN-back | SML | -0.61 | -0.01 | 1.85 | 0.740 | 0.815 |
| Fluid Cognition Composite | EN-back | AUD | -0.59 | -0.01 | 1.94 | 0.762 | 0.815 |
| Fluid Cognition Composite | EN-back | VIS | -5.17 | -0.05 | 2.29 | 0.024 | 0.144 |
| Crystallized Cognition Composite | EN-back | Whole brain | -4.97 | -0.06 | 1.96 | 0.011 | - |
| Crystallized Cognition Composite | EN-back | FPN | -3.13 | -0.05 | 1.46 | 0.032 | 0.107 |
| Crystallized Cognition Composite | EN-back | CON | -4.43 | -0.07 | 1.41 | 0.002 | 0.021 |
| Crystallized Cognition Composite | EN-back | DMN | -2.73 | -0.04 | 1.40 | 0.051 | 0.107 |
| Crystallized Cognition Composite | EN-back | SAL | -1.84 | -0.04 | 1.04 | 0.078 | 0.133 |
| Crystallized Cognition Composite | EN-back | CPN | -1.43 | -0.03 | 1.04 | 0.171 | 0.228 |
| Crystallized Cognition Composite | EN-back | RST | -1.28 | -0.03 | 1.09 | 0.239 | 0.287 |
| Crystallized Cognition Composite | EN-back | DAN | -0.52 | -0.01 | 1.64 | 0.750 | 0.750 |
| Crystallized Cognition Composite | EN-back | VAN | -2.25 | -0.03 | 1.48 | 0.129 | 0.193 |
| Crystallized Cognition Composite | EN-back | SMD | -3.37 | -0.05 | 1.43 | 0.019 | 0.107 |
| Crystallized Cognition Composite | EN-back | SML | -2.26 | -0.04 | 1.17 | 0.054 | 0.107 |
| Crystallized Cognition Composite | EN-back | AUD | -1.03 | -0.02 | 1.23 | 0.403 | 0.439 |
| Crystallized Cognition Composite | EN-back | VIS | -2.94 | -0.05 | 1.45 | 0.042 | 0.107 |

1 EN-back = emotional n-back task, SST = stop signal task, SSRT = stop signal response time, FPN = fronto-parietal, CON = cingulo-opercular, DMN = default mode, SAL = salience, CPN = cingulo-parietal, RST = retrosplenial temporal, DAN = dorsal attention, VAN = ventral attention, SMD = dorsal somatomotor, SML = lateral somatomotor, AUD = auditory, VIS = visual

Conversely, greater fronto-parietal (*β* = 0.04, *SE* = 0.67, adjusted-*p* = .026) and ventral attention (*β* = 0.04, *SE* = 0.62, adjusted-*p* = .016) network functional connectivity similarity between the resting state and the SST related to greater crystallized cognition. All other adjusted-*p* values > .168 (**Table 4**).

**Table 4.**
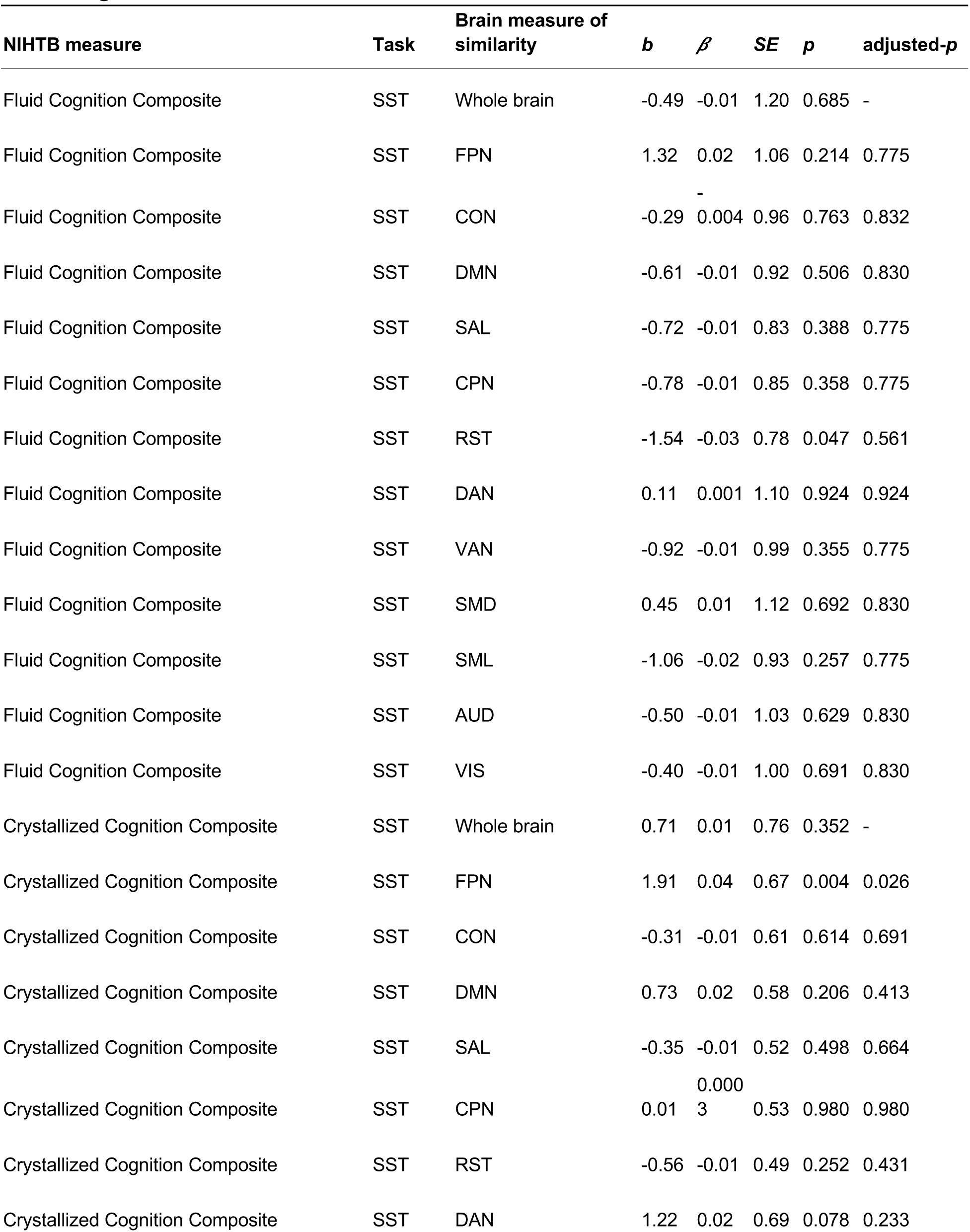
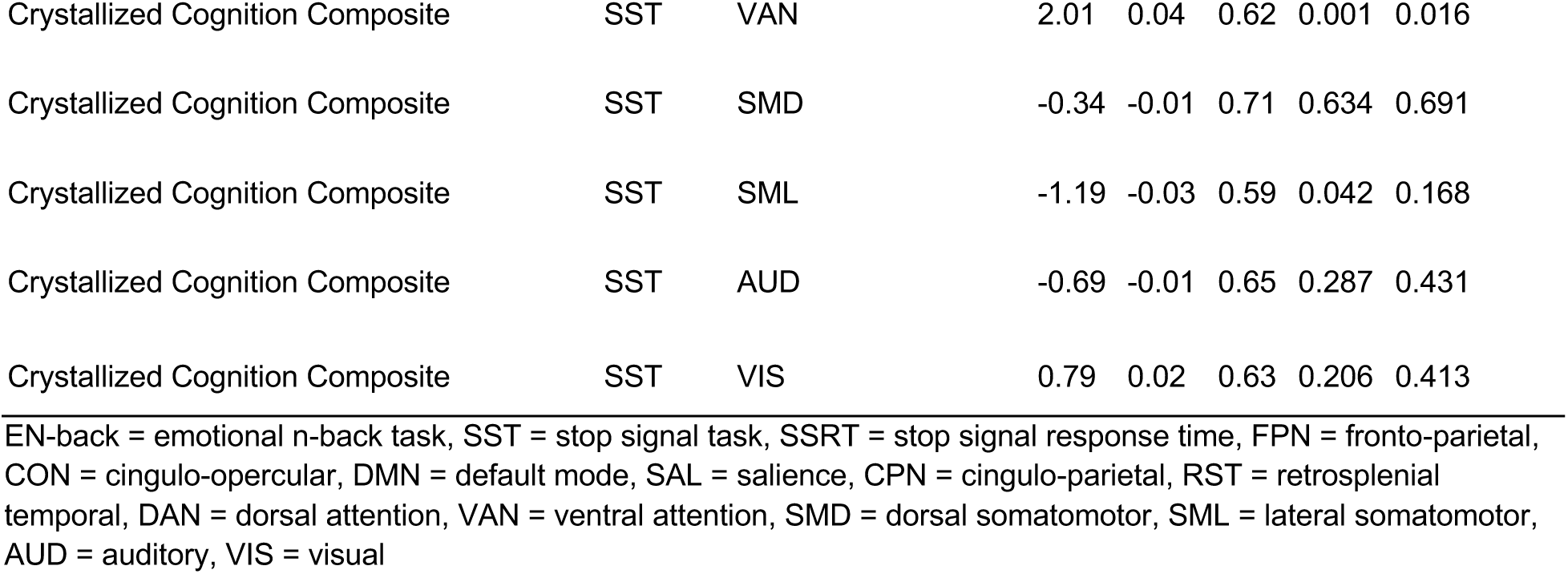
Functional connectivity similarity between the resting state and SST related to broad cognitive measures.

### 3.3. Age-related differences in functional connectivity similarity

Next, we tested if functional connectivity similarity between the resting and task states varied with age (**Figure 3**). We found that functional connectivity similarity between the resting state and the EN-back task did not significantly relate to age (all adjusted-*p* values > .280; **Table 5**). However, functional connectivity similarity between the resting state and the SST of the whole brain (*β* = 0.12, *SE* = 0.0002, *p* < .001), as well as all individual networks (all adjusted-*p* values < .001), increased with age (**Table 5**).

**Figure 3.**
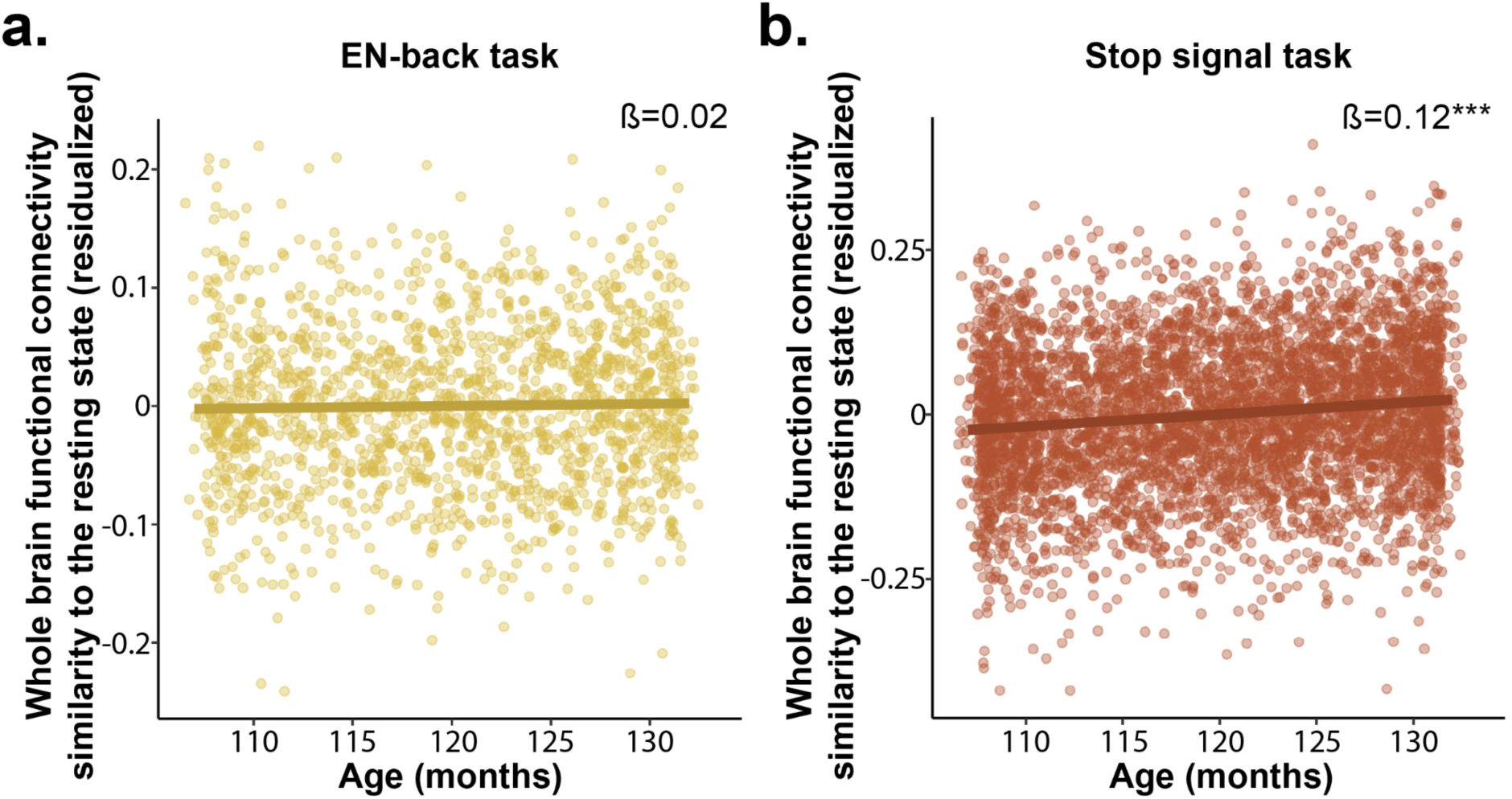
Functional connectivity similarity between the resting state and the SST is related to age. a. Whole brain functional connectivity similarity between the resting state and the EN-back task (yellow) is not significantly related to age. b. Whole brain functional connectivity similarity between the resting state and the SST (red) increases with age. EN-back = Emotional n-back task; SST = Stop signal task; \**p*<.05; \*\**p*<.01; \*\*\**p*<.001

**Table 5.**
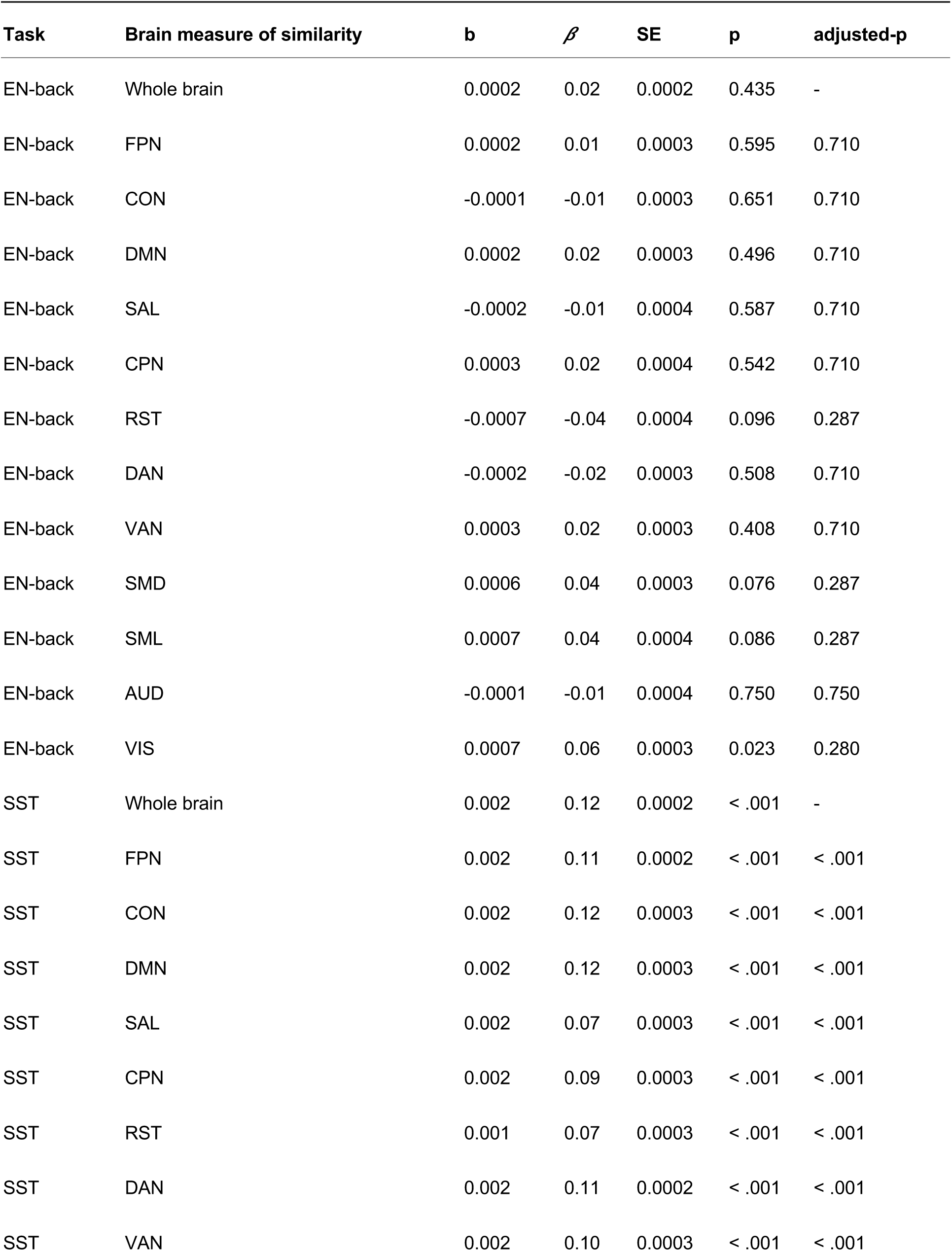
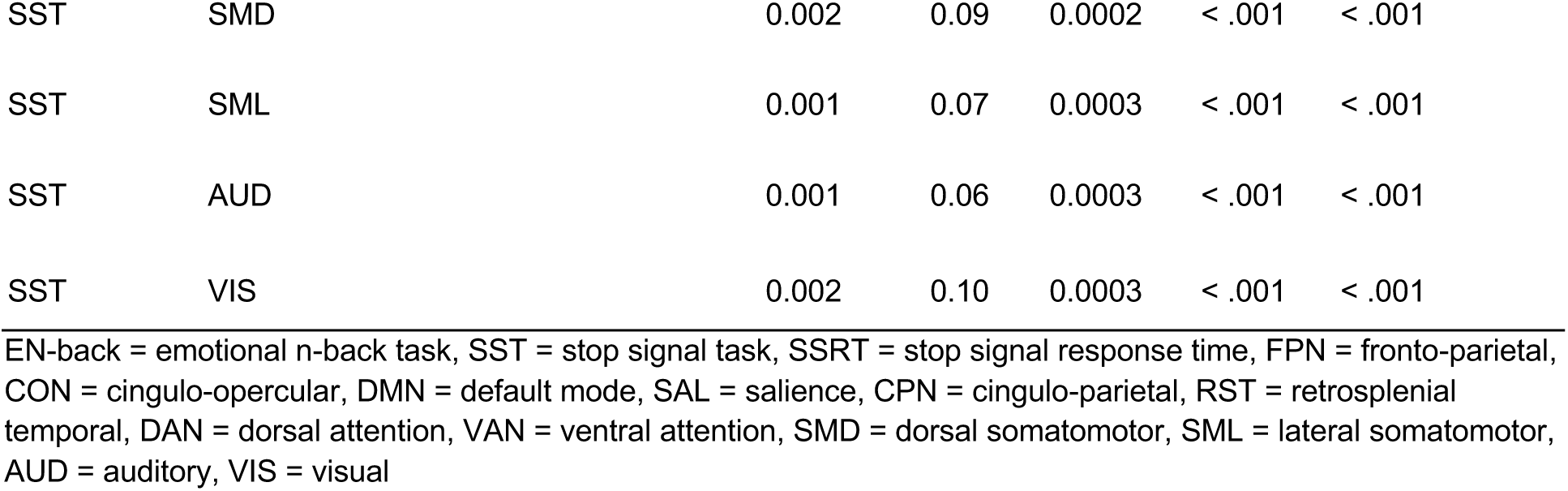
Resting state and task-based functional connectivity similarity related to age.

| Task | Brain measure of similarity | b | $\beta$ | SE | p | adjusted-p |
| --- | --- | --- | --- | --- | --- | --- |
| EN-back | Whole brain | 0.0002 | 0.02 | 0.0002 | 0.435 | - |
| EN-back | FPN | 0.0002 | 0.01 | 0.0003 | 0.595 | 0.710 |
| EN-back | CON | -0.0001 | -0.01 | 0.0003 | 0.651 | 0.710 |
| EN-back | DMN | 0.0002 | 0.02 | 0.0003 | 0.496 | 0.710 |
| EN-back | SAL | -0.0002 | -0.01 | 0.0004 | 0.587 | 0.710 |
| EN-back | CPN | 0.0003 | 0.02 | 0.0004 | 0.542 | 0.710 |
| EN-back | RST | -0.0007 | -0.04 | 0.0004 | 0.096 | 0.287 |
| EN-back | DAN | -0.0002 | -0.02 | 0.0003 | 0.508 | 0.710 |
| EN-back | VAN | 0.0003 | 0.02 | 0.0003 | 0.408 | 0.710 |
| EN-back | SMD | 0.0006 | 0.04 | 0.0003 | 0.076 | 0.287 |
| EN-back | SML | 0.0007 | 0.04 | 0.0004 | 0.086 | 0.287 |
| EN-back | AUD | -0.0001 | -0.01 | 0.0004 | 0.750 | 0.750 |
| EN-back | VIS | 0.0007 | 0.06 | 0.0003 | 0.023 | 0.280 |
| SST | Whole brain | 0.002 | 0.12 | 0.0002 | < .001 | - |
| SST | FPN | 0.002 | 0.11 | 0.0002 | < .001 | < .001 |
| SST | CON | 0.002 | 0.12 | 0.0003 | < .001 | < .001 |
| SST | DMN | 0.002 | 0.12 | 0.0003 | < .001 | < .001 |
| SST | SAL | 0.002 | 0.07 | 0.0003 | < .001 | < .001 |
| SST | CPN | 0.002 | 0.09 | 0.0003 | < .001 | < .001 |
| SST | RST | 0.001 | 0.07 | 0.0003 | < .001 | < .001 |
| SST | DAN | 0.002 | 0.11 | 0.0002 | < .001 | < .001 |
| SST | VAN | 0.002 | 0.10 | 0.0003 | < .001 | < .001 |
| SST | SMD | 0.002 | 0.09 | 0.0002 | < .001 | < .001 |
| SST | SML | 0.001 | 0.07 | 0.0003 | < .001 | < .001 |
| SST | AUD | 0.001 | 0.06 | 0.0003 | < .001 | < .001 |
| SST | VIS | 0.002 | 0.10 | 0.0003 | < .001 | < .001 |

EN-back = emotional n-back task, SST = stop signal task, SSRT = stop signal response time, FPN = fronto-parietal, CON = cingulo-opercular, DMN = default mode, SAL = salience, CPN = cingulo-parietal, RST = retrosplenial temporal, DAN = dorsal attention, VAN = ventral attention, SMD = dorsal somatomotor, SML = lateral somatomotor, AUD = auditory, VIS = visual

### 3.4. Resting-state global system segregation related to functional connectivity similarity, age, and cognition

We were also interested in understanding if functional connectivity similarity between the resting and task states varied with resting-state brain network organization, indexed by global system segregation. We found that greater global system segregation during the resting state related to greater whole brain functional connectivity similarity between the resting state and both tasks (all *p* values < .001), as well as greater functional connectivity similarity in each network (all adjusted-*p* values < .039), except for cingulo-opercular and auditory network functional connectivity similarity between the resting state and the EN-back task (adjusted-*p* values > .112; **Table 6**).

**Table 6.**
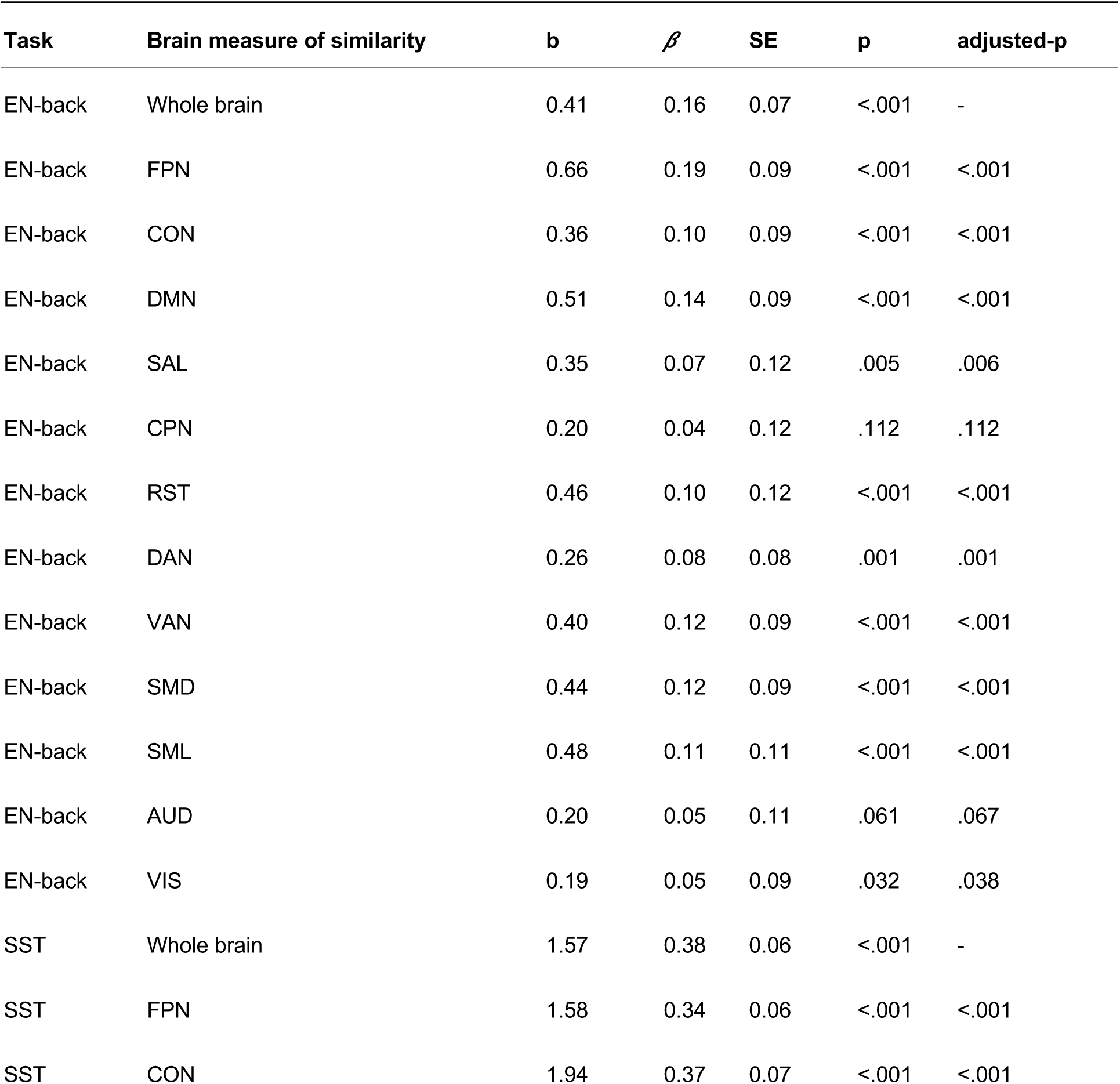
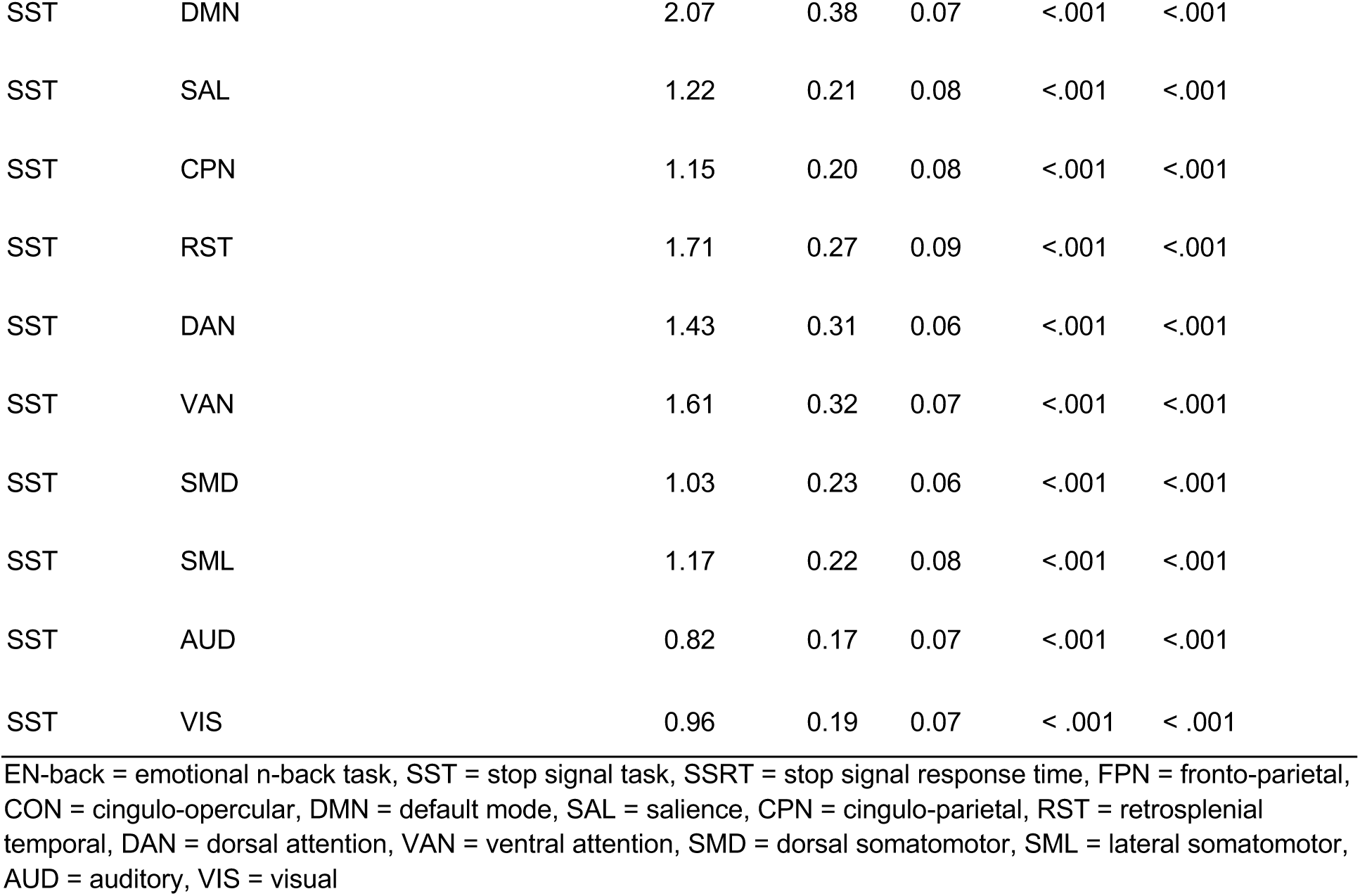
Relationships between global system segregation and functional connectivity similarity.

| Task | Brain measure of similarity | b | $\beta$ | SE | p | adjusted-p |
| --- | --- | --- | --- | --- | --- | --- |
| EN-back | Whole brain | 0.41 | 0.16 | 0.07 | <.001 | - |
| EN-back | FPN | 0.66 | 0.19 | 0.09 | <.001 | <.001 |
| EN-back | CON | 0.36 | 0.10 | 0.09 | <.001 | <.001 |
| EN-back | DMN | 0.51 | 0.14 | 0.09 | <.001 | <.001 |
| EN-back | SAL | 0.35 | 0.07 | 0.12 | .005 | .006 |
| EN-back | CPN | 0.20 | 0.04 | 0.12 | .112 | .112 |
| EN-back | RST | 0.46 | 0.10 | 0.12 | <.001 | <.001 |
| EN-back | DAN | 0.26 | 0.08 | 0.08 | .001 | .001 |
| EN-back | VAN | 0.40 | 0.12 | 0.09 | <.001 | <.001 |
| EN-back | SMD | 0.44 | 0.12 | 0.09 | <.001 | <.001 |
| EN-back | SML | 0.48 | 0.11 | 0.11 | <.001 | <.001 |
| EN-back | AUD | 0.20 | 0.05 | 0.11 | .061 | .067 |
| EN-back | VIS | 0.19 | 0.05 | 0.09 | .032 | .038 |
| SST | Whole brain | 1.57 | 0.38 | 0.06 | <.001 | - |
| SST | FPN | 1.58 | 0.34 | 0.06 | <.001 | <.001 |
| SST | CON | 1.94 | 0.37 | 0.07 | <.001 | <.001 |
| SST | DMN | 2.07 | 0.38 | 0.07 | <.001 | <.001 |
| SST | SAL | 1.22 | 0.21 | 0.08 | <.001 | <.001 |
| SST | CPN | 1.15 | 0.20 | 0.08 | <.001 | <.001 |
| SST | RST | 1.71 | 0.27 | 0.09 | <.001 | <.001 |
| SST | DAN | 1.43 | 0.31 | 0.06 | <.001 | <.001 |
| SST | VAN | 1.61 | 0.32 | 0.07 | <.001 | <.001 |
| SST | SMD | 1.03 | 0.23 | 0.06 | <.001 | <.001 |
| SST | SML | 1.17 | 0.22 | 0.08 | <.001 | <.001 |
| SST | AUD | 0.82 | 0.17 | 0.07 | <.001 | <.001 |
| SST | VIS | 0.96 | 0.19 | 0.07 | < .001 | < .001 |

1 EN-back = emotional n-back task, SST = stop signal task, SSRT = stop signal response time, FPN = fronto-parietal, CON = cingulo-opercular, DMN = default mode, SAL = salience, CPN = cingulo-parietal, RST = retrosplenial temporal, DAN = dorsal attention, VAN = ventral attention, SMD = dorsal somatomotor, SML = lateral somatomotor, AUD = auditory, VIS = visual

Given prior evidence that brain network segregation increases across childhood and adolescence (Sun et al., 2024, 2024; Tooley, Park, et al., 2022) and relates to cognitive control (Chan et al., 2014; Keller et al., 2023; Kong et al., 2020), we sought to test if this effect was present in our limited age range. Indeed, global system segregation increased across age (*β* = 0.03, *SE* = 5.18e-05, *p* = .018). Additionally greater global system segregation was related to greater 2-back accuracy (*β* = 0.05, *SE* = 0.04, adjusted-*p* = .003), but not significantly related to SSRT (adjusted-*p* = .660), fluid cognition composite (adjusted-*p* = .105), nor crystallized cognition composite (adjusted-*p* = .894).

To understand if the relationships between functional connectivity and cognition could be driven by global system segregation, we tested if system segregation accounted for the relationships we identified between functional connectivity similarity and cognition. We found that even when controlling for system segregation, all previously identified relationships between functional connectivity similarity and cognition remained significant (all adjusted-*p* values < .013; **Table 7**).

**Table 7.** Functional connectivity similarity related to task performance and broad cognitive measures after controlling for global system segregation.

| Cognitive measure | Task | Brain measure of similarity | *b* | $\beta$ | *SE* | *p* | adjusted-*p* |
| --- | --- | --- | --- | --- | --- | --- | --- |
| 2-back accuracy | EN-back | Whole brain | -0.09 | -0.08 | 0.03 | .001 | .002 |
| 2-back accuracy | EN-back | CON | -0.13 | -0.15 | 0.02 | <.001 | <.001 |
| 2-back accuracy | EN-back | SAL | -0.06 | -0.09 | 0.02 | <.001 | <.001 |
| 2-back accuracy | EN-back | SMD | -0.09 | -0.11 | 0.02 | <.001 | <.001 |
| SSRT | SST | Whole brain | 30.05 | 0.06 | 8.93 | .001 | .002 |
| SSRT | SST | FPN | 26.50 | 0.05 | 7.73 | .001 | .002 |
| SSRT | SST | DMN | 32.78 | 0.08 | 6.83 | <.001 | <.001 |
| SSRT | SST | CPN | 16.63 | 0.04 | 5.92 | .005 | .006 |
| SSRT | SST | DAN | 31.15 | 0.06 | 7.98 | <.001 | <.001 |
| SSRT | SST | VAN | 20.88 | 0.05 | 7.21 | .004 | .005 |
| SSRT | SST | SMD | 24.01 | 0.05 | 7.92 | .002 | .003 |
| Fluid Cognition Composite | EN-back | Whole brain | -7.80 | -0.06 | 3.15 | .013 | .013 |
| Crystallized Cognition Composite | EN-back | Whole brain | -5.48 | -0.06 | 1.98 | .006 | .006 |
| Crystallized Cognition Composite | EN-back | FPN | -4.65 | -0.07 | 1.42 | .001 | .002 |
| Crystallized Cognition Composite | SST | CON | 2.16 | 0.04 | 0.71 | .002 | .003 |
| Crystallized Cognition Composite | SST | VAN | 2.25 | 0.05 | 0.66 | .001 | .002 |

EN-back = emotional n-back task, SST = stop signal task, SSRT = stop signal response time, FPN = fronto-parietal, CON = cingulo-opercular, DMN = default mode, SAL = salience, CPN = cingulo-parietal, DAN = dorsal attention, VAN = ventral attention, SMD = dorsal somatomotor

## 4. Discussion

Here, we examined how cognition in youth is supported by reconfiguration of functional brain network architecture in response to changing cognitive demands. To do this, we operationalized reconfiguration by calculating network similarity between the resting state and two executive function tasks, tapping working memory and response inhibition. This measure of functional connectivity similarity is the opposite of reconfiguration, such that as similarity increases, reconfiguration decreases. For clarity and to align with previous work, we will use the term reconfiguration to discuss our findings. We tested if the degree of reconfiguration was related to executive function performance on the tasks done in the MRI, as well as to broad measures of fluid and crystallized cognition. In adults, there is evidence that less reconfiguration of functional brain network organization across various cognitive states is beneficial for performance on language, reasoning, and working memory tasks, as well as general intelligence (Schultz & Cole, 2016). Schultz and Cole (2016) suggest that an intrinsic functional brain network organization that is “preconfigured” to facilitate a variety of cognitive processes with only minute adjustments for different cognitive processes enables maximal efficiency. In late childhood though, we find an opposite relationship, such that functional network reconfiguration, or lower values of our similarity metric, between the resting state and cognitively-demanding tasks is beneficial for cognition, while a preconfigured intrinsic network organization is detrimental. We found evidence that more reconfiguration between the resting state and both tasks predicts better task performance and, in the case of reconfiguration between the resting state and working memory, better fluid and crystallized cognition. Importantly, these relationships are seen above and beyond relationships with resting-state brain network segregation, a marker of brain maturation (Keller et al., 2023).

We conceptualize the relationship between the degree of functional brain network reconfiguration when faced with cognitive demands and cognitive performance across a U-shaped spectrum. One extreme end of the spectrum is characterized by rigidity, or a principal functional network organization that resists adapting to new demands. The other end is characterized by disorganization, where every task requires an organizational overhaul. Neither end of this spectrum is particularly efficient or beneficial for cognition and there is likely a balance point along this spectrum that maximizes efficiency and supports a variety of cognitive demands. Our findings, in conjunction with evidence in adults (Schultz & Cole, 2016), suggest that adults and children differ in their optimal balance point along this spectrum, with optimal organizational arrangement in adults favoring more preconfiguration, toward the rigidity end, and optimal arrangement in late childhood favoring more flexibility, toward the disorganization end. The discrepancy in the optimal functional network reconfiguration between adults and children gives rise to the question: *Why is it beneficial to be less preconfigured, or more flexible, in late childhood?*

Efficient preconfiguration requires a functional network organization that is both stable (present both at the resting state and during tasks) and utilitarian (supports a variety of cognitive demands). A well-established and stable brain organization may be the goal, but is not the reality in the transition between childhood and adolescence. Late childhood into adolescence is a time of marked brain plasticity (Fuhrmann et al., 2015). This second wave of plasticity presents a unique window of time wherein cortical sensitivity to aspects of the environment facilitates cognitive maturation (Larsen & Luna, 2018). During these windows, the same neural mechanisms driving this period of plasticity may be those that are staving off preconfiguration. For example, during late childhood the spatial boundaries of functional brain networks are not fully established and show subtle refinements throughout adolescence (Cui et al., 2020; Marek et al., 2015; Sun et al., 2024; Tooley, Bassett, et al., 2022). Throughout adolescence local homogeneity of functional brain networks decreases, further speaking to refinement and network specialization continuing across this period (Calabro et al., 2025). Additionally, cortical microstructure is still undergoing significant change, particularly in transmodal association cortex (Baum et al., 2022). Preconfiguration during such a period of change could place undue and premature restraint on the refinement of brain organization. Thus, a lack of preconfiguration during late childhood and adolescence could necessitate larger changes to transition from the resting state to successfully performing a task.

As the brain is maturing, resting-state functional brain networks are becoming more distinct from each other, or segregated (Keller et al., 2023; Sun et al., 2024; Tooley, Park, et al., 2022), which we see reflected in our sample despite the restricted age range. Still, in this age range, we see flexibility and a lack of preconfiguration is related to improved cognition, even in those individuals who have more segregated brain networks during the resting-state. This speaks to a qualitative difference between childhood and adulthood, wherein more reconfiguration relates to better cognition. We posit that the degree of change from the resting state to a cognitively-demanding task state reflects flexibility driven by developmentally-appropriate plasticity in contrast to the preconfigured efficiency of the adult brain.

We noted some interesting differences between tasks which suggest that the utility of preconfiguration likely differs between cognitive domains. Regarding the working memory task, more reconfiguration was related to better crystallized and fluid cognition. This mirrors the pattern of results relating reconfiguration to performance on the working memory task. This pattern emerged when looking at reconfiguration across the whole brain and specifically at the cingulo-opercular network. Whole brain reconfiguration between the resting state and the inhibition task, however, was not related to any broad measures of cognition. Only reconfiguration in the fronto-parietal and ventral attention networks were related to crystalized cognition and this relationship was opposite what was seen for working memory - greater reconfiguration related to worse cognition. Taken together, we see in late childhood the relationship between reconfiguration and broad cognitive measures is dependent on the cognitive domain, which is not the case in adult work where the relationship between reconfiguration and intelligence is the same across cognitive domains (Schultz & Cole, 2016). Additionally, reconfiguration between the resting state and the response inhibition task, but not the working memory task, increased with age. In adolescence, reconfiguration from the resting state to other tasks decreases with age and has been linked with early life stress (Lee et al., 2024). Examining reconfiguration over a wider age range and across additional cognitive contexts will undoubtedly help unpack these nuances and the ABCD Study can support these investigations in the future.

To further understand developmental trajectories of reconfiguration, it is useful to consider the mechanisms that support brain change based on task demands. One candidate mechanism that may drive changes in reconfiguration is the development of highly integrated functional zones, also called “hubs”. Hubs are areas of the brain characterized by strong patterns of communication both locally within a network and globally across networks (Gratton et al., 2018) and thereby facilitate efficient communication throughout the brain (Cole et al., 2013; van den Heuvel & Sporns, 2013). Hub activation and connectivity have been linked to cognition (Bertolero et al., 2018; Betzel et al., 2014; Warren et al., 2014), and in adults hubs exhibit more flexible functional connectivity across tasks compared to other brain regions (Gratton et al., 2016). The development of functional hubs can be conceptualized as the brain’s maturing switchboard for coordinating large-scale network communications. Early hub function is constrained by immature long-range fibers, which undergo myelination into adulthood (van den Heuvel et al., 2018). As these fibers mature through childhood and adolescence, functional hubs shift from sensorimotor to association areas (Oldham & Fornito, 2019) and coordinate connectivity among association networks, as well as between association and sensory networks (Demeter et al., 2023). Hubs exhibiting strong functional connectivity across association and sensory networks support cognitive control in childhood and adolescence (Demeter et al., 2023). We posit hubs as a central driver of the reconfiguration required for different tasks demands, facilitating contextual changes in connectivity across established brain networks. Future work that closely examines developmental trajectories of myelination and hub connectivity across task contexts will help clarify how, when, and under what conditions preconfiguration becomes more advantageous as the brain matures into adulthood.

## 5. Conclusion

A functional brain network organization primed to carry out many tasks with only a few small organizational tweaks is beneficial in adulthood. Here, we asked if this relationship held true in children. We found that in late childhood more reconfiguration, or a larger brain network organizational overhaul, to carry out a cognitively-demanding task, was related to better performance on the task at hand and, in some cases, generalized to more wide-ranging measures of cognition. The contrast of our findings with prior work in adults highlights a qualitative developmental shift along a spectrum from flexibility providing a cognitive advantage in childhood to preconfiguration providing a cognitive advantage in adulthood. Reconfiguration also decreases with age, suggesting there may be a developmental pattern of flexibility early in life giving way to preconfiguration that emerges with maturation and unfolds at unique timescales for different cognitive domains.

## Data Statement

Data used in the preparation of this article were obtained from the Adolescent Brain Cognitive Development^SM^ (ABCD) Study (https://abcdstudy.org), held in the NIMH Data Archive (NDA). The ABCD data used in this report came from NDA Release 3.0 (DOI: http://dx.doi.org/10.15154/1520591).

## Author contributions: CRediT

MEM - Conceptualization, Methodology, Formal analysis, Visualization, Writing – original draft, Writing – review and editing

AJJ - Writing – original draft, Writing – review and editing

TN - Conceptualization, Methodology, Writing – original draft, Writing – review and editing

The authors declare no conflicts of interest.

