## Supplemental Materials for "Cognition is associated with task-related brain network reconfiguration in late childhood"

**Supplementary Materials**

*Differences in functional connectivity similarity between the resting state and two executive function tasks*

We additionally tested for differences in functional connectivity similarity between the resting state and the EN-back task and the resting state and the SST. To do this we estimated linear mixed effects models, each with the one functional connectivity similarity value as the outcome and a categorical variable labeling the comparison (i.e., resting state-SST, resting state-EN-back) as the predictor of interest. We included covariates of age, sex, socioeconomic status composite score, race/ethnicity, and parent marital status, as well as a random intercept of participant. We corrected for multiple comparisons among the p-values for the 12 network models using a FDR correction at $q < .05$. Additionally, we ran an F test to compare the two variances in functional connectivity similarity between the resting state and the EN-back task and between the resting state and the SST.

We found that whole brain functional connectivity similarity between the resting state and the SST task was significantly greater than similarity between the resting state and the EN-back task ($\beta = 1.69$, $SE = 0.002$, $p < .001$; **Supplementary Table 1**; **Supplementary Figure 1**). Further, functional connectivity similarity between the resting state and the EN-back task was significantly less than similarity between the resting state and the SST in each network (all adjusted-$p$ values < .001; **Supplementary Table 1**; **Supplementary Figure 2**).

Additionally, the variance in functional connectivity similarity between the resting state and the SST was significantly greater than variance in functional connectivity similarity between the resting state and the EN-back task ($F(4677, 1594) = 2.65$, $p < .001$).

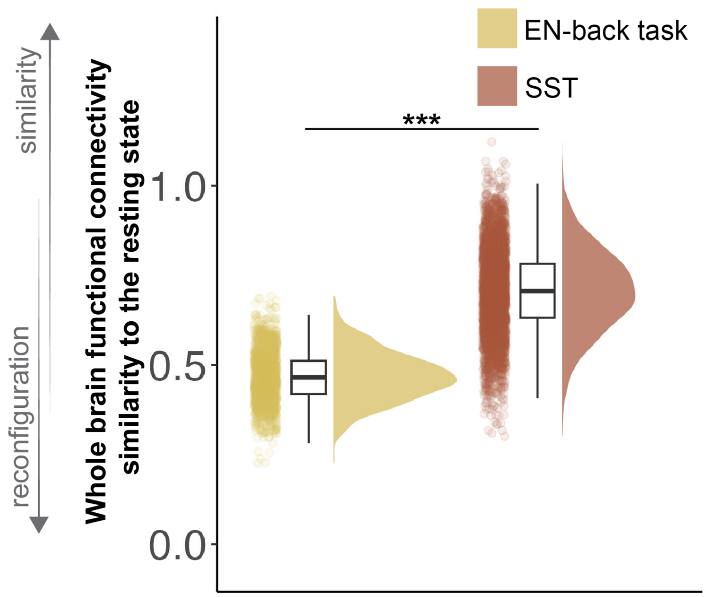


**Supplementary Figure 1. Whole brain functional connectivity similarity to the resting state is greater in the SST compared to the EN-back task.** *$p$<.05; **$p$<.01; ***$p$<.001

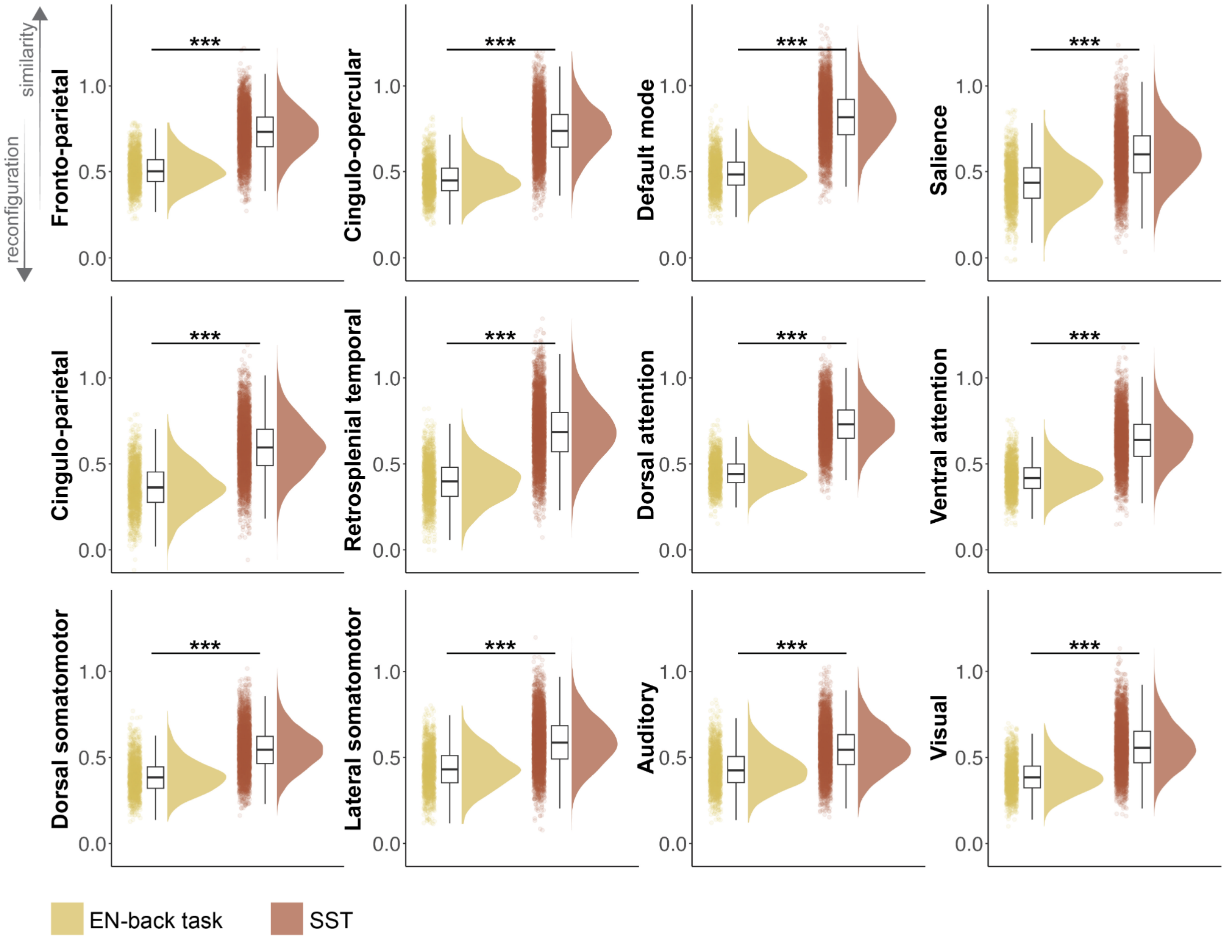


**Supplementary Figure 2. Network functional connectivity similarity to the resting state is greater in the SST compared to the EN-back task.** Y-axes are resting state functional connectivity similarity for each network in the Gordon 2016 parcellation (Gordon et al., 2016) with regard to the EN-back task (yellow) and the SST (red). EN-back = Emotional n-back task; SST = Stop signal task; **p*<.05; ***p*<.01; ****p*<.001

**Supplementary Table 1. Differences in functional connectivity similarity between the resting state and two executive function tasks**

| **Brain measure of similarity** | ***b*** | ***β*** | ***SE*** | ***p*** | **adjusted-*p*** |
|---|---|---|---|---|---|
| Whole brain | 0.25 | 1.69 | 0.002 | < .001 | - |
| FPN | 0.23 | 1.50 | 0.003 | < .001 | < .001 |
| CON | 0.29 | 1.61 | 0.003 | < .001 | < .001 |
| DMN | 0.34 | 1.70 | 0.003 | < .001 | < .001 |
| SAL | 0.17 | 1.03 | 0.004 | < .001 | < .001 |
| CPN | 0.24 | 1.29 | 0.004 | < .001 | < .001 |
| RST | 0.30 | 1.45 | 0.004 | < .001 | < .001 |
| DAN | 0.29 | 1.72 | 0.003 | < .001 | < .001 |
| VAN | 0.23 | 1.44 | 0.003 | < .001 | < .001 |
| SMD | 0.16 | 1.23 | 0.003 | < .001 | < .001 |
| SML | 0.16 | 1.05 | 0.003 | < .001 | < .001 |
| AUD | 0.12 | 0.92 | 0.003 | < .001 | < .001 |
| VIS | 0.18 | 1.23 | 0.003 | < .001 | < .001 |

FPN = fronto-parietal, CON = cingulo-opercular, DMN = default mode, SAL = salience, CPN = cingulo-parietal, RST = retrosplenial temporal, DAN = dorsal attention, VAN = ventral attention, SMD = dorsal somatomotor, SML = lateral somatomotor, AUD = auditory, VIS = visual
